# All-optical spatiotemporal mapping of ROS dynamics across mitochondrial microdomains *in situ*

**DOI:** 10.1101/2023.01.07.523093

**Authors:** Shon A. Koren, Nada A. Selim, Lizbeth De La Rosa, Jacob Horn, M. Arsalan Farooqi, Alicia Y. Wei, Annika Müller-Eigner, Jacen Emerson, Gail V.W. Johnson, Andrew P. Wojtovich

**Affiliations:** University of Rochester Medical Center, Department of Anesthesiology and Perioperative Medicine, 575 Elmwood Ave., Rochester NY, 14642 Box 711/604. United States of America; University of Rochester Medical Center, Department of Pharmacology and Physiology, 575 Elmwood Ave., Rochester NY, 14642 Box 711/604. United States of America; Research Group Epigenetics, Metabolism and Longevity, Research Institute for Farm Animal Biology (FBN), Dummerstorf, 18196, Germany

## Abstract

Hydrogen peroxide (H_2_O_2_) functions as a second messenger to signal metabolic distress through highly compartmentalized production in mitochondria. The dynamics of ROS generation and diffusion between mitochondrial compartments and into the cytosol govern oxidative stress responses and pathology, though our understanding of these processes remains limited. Here, we couple the H_2_O_2_ biosensor, HyPer7, with optogenetic stimulation of the ROS-generating protein KillerRed targeted into multiple mitochondrial microdomains. Single mitochondrial photogeneration of H_2_O_2_ demonstrates the spatiotemporal dynamics of ROS diffusion and transient hyperfusion of mitochondria due to ROS. Measurement of microdomain-specific H_2_O_2_ diffusion kinetics reveals directionally selective diffusion through mitochondrial microdomains. All-optical generation and detection of physiologically-relevant concentrations of H_2_O_2_ between mitochondrial compartments provide a map of mitochondrial H_2_O_2_ diffusion dynamics *in situ*. These kinetic details of spatiotemporal ROS dynamics and inter-mitochondrial spreading forms a framework to understand the role of ROS in health and disease.

## Introduction

Mitochondrial reactive oxygen species (ROS) act as metabolic signals to coordinate bioenergetic state with cellular activity and adaptation. Under normal metabolic activity, the electron transport chain (ETC) produces low levels of ROS in the form of superoxide (O_2_^−^) which rapidly converts into hydrogen peroxide (H_2_O_2_) through dismutation^1^. Since H_2_O_2_ readily crosses membranes, it can act as a second messenger akin to Ca^2+^. ROS accumulates at sub-toxic levels to regulate metabolic signaling. This basal redox tone is defined as oxidative eustress^2–4^. Pathological conditions increase ROS production in mitochondria beyond this basal state, termed oxidative distress. In disease, H_2_O_2_ accumulates and is released from mitochondria to aberrantly oxidize and impair cellular machinery, ultimately inducing apoptosis^5^.

Years of investigating oxidative stress in disease has emphasized the role of ROS in driving cellular pathology^6–8^, but surprisingly little is known about the “how” and “when” ROS generated in mitochondria drive global cellular changes. The exact concentration difference between healthy, metabolically signaling ROS and oxidative damage remains unknown. Once generated, ROS can diffuse between mitochondrial compartments and into the cytosol, but the kinetics of this process is only beginning to be characterized^9,10^. Moreover, the four-decade old hypothesis^11^ that ROS releases from the mitochondrial matrix into the cytosol has only minimal experimental evidence^12^. Deeper understanding of these processes would elucidate how ROS coordinates metabolic signaling and oxidative stress pathways.

Mitochondrial H_2_O_2_ generation and diffusion is highly spatially organized and complex. Multiple sites along the ETC generate ROS at different rates, thereby forming distinct microdomains of H_2_O_2_ dynamics in mitochondria such as the matrix, inter- and outer-membrane spaces (IMS and OMS, respectively)^1,13–15^. Mitochondrial microdomains differ in the selectivity and rate of ROS scavenging^16^, the propensity of H_2_O_2_ to diffuse due to distinct membrane porosity^17–20^, and protein substrate crowding^21–23^, which can alter downstream ROS signal propagation. Individual sites of ROS production in mitochondrial microdomains are linked to distinct diseases which further highlights the variation in local ROS production^6^. Beyond this biological complexity, methods to induce and measure ROS production have been historically restricted to drug treatments or genetic manipulation. These methods provide either spatial or temporal information, but not both. Recent advances have bypassed these constraints with targeted expression of H_2_O_2_-generating proteins such as the D-alanine oxidase (DAO) system^24^. These systems provide a high degree of spatial control of H_2_O_2_ production but at relatively slow timescales and with limited temporal regulation^9,12^. Furthermore, by producing H_2_O_2_ and not superoxide to mimic ROS generation at the ETC, these systems may not fully recapitulate kinetic details of ROS diffusion dynamics.

Here, we employ an all-optical ROS generation and detection system to provide spatial and temporal control with endogenous dismutation in living cells. We use the superoxide ROS-generating protein tandem-KillerRed (KR)^25–27^ and the H_2_O_2_ biosensor HyPer7^28^ to measure ROS dynamics throughout mitochondrial microdomains. Spatially restricted ROS generation and monitoring in single mitochondria demonstrates the kinetics of ROS diffusion across subpopulations of mitochondria in a cell alongside the transient hyperfusion of mitochondria. Controlling for HyPer7 biosensor characteristics in each mitochondrial microdomain reveals that ROS exhibits directionally selective diffusion into the matrix. Using this all-optical ROS generation and detection technique, we systematically generate a map of ROS diffusion kinetics into and out of each mitochondrial microdomain in living cells. We record the release of H_2_O_2_ from the matrix of single mitochondria into the cytosol and into surrounding mitochondria about one μm away in normal metabolic conditions. Together, these data suggest a model of how ROS generated in mitochondrial compartments can spread throughout a cell on the minute timescale.

## Results

### Optical control of ROS generation in single mitochondria in situ

ROS diffusion between mitochondria and the cytosol governs cellular responses to changes in metabolism, but the dynamics of ROS movement between these regions remains unknown. To monitor ROS dynamics in mitochondria of living cells, we first co-expressed the superoxide-generating protein tandem-KillerRed (KR) and H_2_O_2_ biosensor HyPer7, both targeted into the matrix of mitochondria in cells. The photosensitizer KR has a known superoxide quantum yield^29^. HyPer7 has been previously shown to be highly selective for H_2_O_2_ and sensitive enough to detect gradients of mitochondrial ROS in living organisms^12,28,30^. We reasoned that coupling expression of these two proteins could record single mitochondrial ROS responses **(Fig. 1A**).

**Figure 1:**
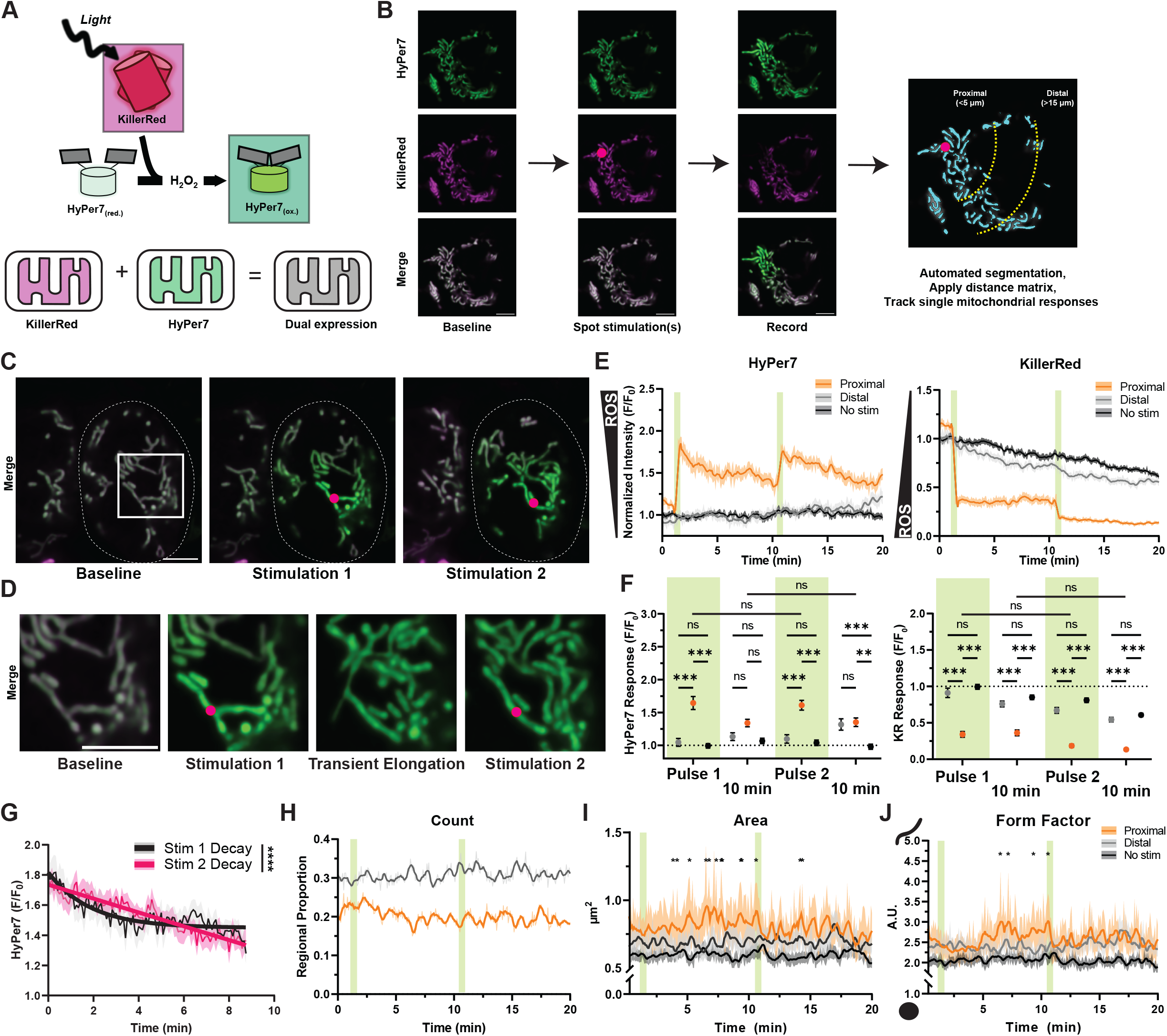
Spatiotemporal manipulation and monitoring of ROS in single mitochondria. A. Schematic representation of all optical ROS generation and detection in mitochondria B. Imaging and analysis workflow to track spatiotemporal mitochondrial responses relative to single mitochondrial ROS photogeneration. C. Representative images, where dotted line indicates a single cell expressing matrix-targeted HyPer7 and KR. Denoted circle indicates static spot stimulation used to pulse ROS photogeneration. D. Inset from c, highlighting transient elongation and contact of photostimulated mitochondria. E. Single mitochondrial responses of HyPer7 and KR intensities normalized to area and baseline, mean ± SEM, N = 39 distal, 54 proximal, and 95 non-stimulated mitochondria per frame per group, on average. F. Comparison of HyPer7 and KR responses at first frame immediately following pulse 1, 10 minutes following first stimulation, immediately following pulse 2, and 10 minutes following second stimulation. Two-way ANOVA with Tukey *post-hoc* multiple comparisons. G. Decay of HyPer7 following stimulation 1 and 2 fitted with a nonlinear variable plateau followed by single phase decay, mean ± SEM. Decay compared with extra sum-of-squares F test. H. Mitochondrial regional proportion of proximal and distal subpopulations relative to total in the cell over time. I. Area of individual mitochondria in subpopulations over time through ROS photostimulation. Two-way ANOVA with Tukey *post-hoc* multiple comparisons, mean ± SEM. J. Form factor of individual mitochondria in subpopulations over time through ROS photostimulation. Two-way ANOVA with Tukey *post-hoc* multiple comparisons, mean ± SEM. Green bars indicate single frames of KR photostimulation. *p < 0.05, **p < 0.01, ***p < 0.001. Scalebars denote 5 μm.

Single mitochondria in cells expressing matrix-targeted KR and HyPer7 were selectively photostimulated with 5 sec pulses of 561 nm light (88 μW). The HyPer7 and KR intensities of mitochondria in the cell were measured before and after photostimulation to track the diffusion of ROS between mitochondria over time. To account for mitochondrial movement across three-dimensional space, as well as fusion and fission dynamics, we developed a method to automatically segment and track single mitochondria for each frame of 20+ minute recordings (5 sec between frames, >220 frames per experiment). The HyPer7 and KR intensity of each mitochondrion was normalized to its area in that frame to account for changes in area over time. Additionally, the distance of each mitochondrion to the fixed spot stimulation point was calculated so that ROS diffusion could be measured in proximal and distal mitochondrial populations (approximately 5 μm and 15 μm away from spot stimulation, respectively). Together, this system allowed for the spatiotemporal monitoring of single mitochondrial ROS and morphological responses while limiting the effect of mitochondrial drift and size changes (**Fig. 1B**).

Spot photostimulation of KR led to distinct spatiotemporal ROS responses in single mitochondria as measured by HyPer7 (**Fig. 1C-D**). As expected, photostimulating KR led to rapid photobleaching as its chromophore oxidized as a product of photosensitization and ROS generation^25,31,32^. Mitochondria proximal to the spot stimulation had strongly photobleached KR and increased HyPer7 intensity, whereas mitochondria in the same cell which were distal to spot stimulation had a lower response of KR and HyPer7 (**Supplemental Video 1**). Non-stimulated control mitochondria exhibited comparably similar responses to imaging as distal mitochondria (**Supplemental Video 2**), indicating photostimulation of KR was constrained to mitochondria only proximal to the stimulation point. This suggested this technique could detect mitochondrial subpopulation responses to pulsed ROS generation.

Quantifying single mitochondrial subpopulation responses to ROS generation revealed an immediate, roughly 65% increase in HyPer7 intensity in mitochondria proximal to photostimulation (**Fig. 1E-F**). This indicated rapid dismutation of KR-generated superoxide into H_2_O_2_ and subsequent oxidation of the HyPer7 probe^12,28^. After the second photostimulation pulse on KR, proximal mitochondria were repeatedly oxidized to the same level compared to the first pulse despite HyPer7 not being fully reduced back toward baseline, indicating a lower photosensitization capacity for KR following initial photostimulation. This double photostimulation protocol began overwhelming matrix antioxidant machinery, as evinced by the decay kinetics of HyPer7 slowing after the second pulse (K_first_ = 0.54 ± 0.07, K_second_ = 0.085 ± 0.031 in F/F_0_ per sec) (**Fig. 1G**) without a change in final HyPer7 intensity (**Fig. 1F**). Together, these data demonstrated that this all-optical technique of pulse generating H_2_O_2_ reliably modeled oxidative distress with spatiotemporal precision.

### Tracking intracellular spread of spatially restricted mitochondrial ROS

To assess whether matrix-generated ROS could spread between mitochondria in a cell, we measured the oxidation of mitochondrial subpopulations proximal and distal to the photostimulation point. Surprisingly, we measured an increase in oxidation of distal mitochondria after a nearly five-minute delay following the second KR photostimulation pulse (roughly 15 min following the initial ROS pulse). This distal ROS spread correlated with rapid trafficking of oxidized mitochondria to distal regions of the cell (**Supplemental Movie 3**). This trafficking often occurred after a >5 min delay following the second stimulation pulse. This delayed increase in distal mitochondrial oxidation and trafficking was consistent with the idea that the second pulse of KR began overwhelming mitochondrial antioxidant machinery to subsequently impact the local redox environment outside mitochondria.

To probe this inter-mitochondrial diffusion of ROS further, we analyzed mitochondrial morphology before and after ROS photogeneration. Photostimulated mitochondria appeared to briefly elongate and increase contact with neighboring mitochondria (**Fig. 1D, Supplemental Movie 4**). Frame-by-frame analysis of mitochondria revealed relatively stable number of mitochondria in proximal and distal regions measured as a proportion to total (**Fig. 1H**). Contrastingly, mitochondria proximal to photostimulation exhibited transient waves of hyperfusion as measured by area and form factor (**Fig. 1I-J**), a metric which reports the complexity of mitochondrial shape. These waves of mitochondrial elongation continued throughout the duration of recording following an initial delay after the first ROS pulse, but slowed in frequency after the second pulse where distal mitochondria began similarly transiently increasing in area and form factor. Non-stimulated control mitochondria did not undergo measurable changes in morphology, further supporting that these changes were due to spatially restricted ROS generation through KR photostimulation.

These results were largely recapitulated in photostimulation experiments in mouse embryonic fibroblasts (MEFs) co-expressing matrix-targeted HyPer7 and KR (**Extended Fig. 1A-B**). Spot stimulation did minimally oxidize some distal mitochondria which fully decayed to baseline after both pulses of KR photostimulation. MEFs exhibited altered decay kinetics between ROS photogeneration pulses with a faster decay in the second pulse relative to the first (K_first_ = 0.24 ± 0.31, K_second_ = 1.73 ± 0.34 in F/F_0_ per sec) without a difference in final HyPer7 intensity (**Extended Fig. 1C-D**). This ROS photogeneration in MEFs also led to transient mitochondrial hyperfusion as measured by area and form factor (**Extended Fig. 1E**). Together, these results suggested ROS transiently increases mitochondrial fusion, but the intra-cellular spread and decay of ROS may differ between cell types.

### Mitochondrial microdomain-specific responses to exogenous H_2_O_2_

The ROS landscape varies greatly between mitochondrial compartments^1,13,14^. To understand ROS dynamics at each mitochondrial microdomain, we first confirmed proper microdomain-specific targeting of HyPer7. We targeted HyPer7 to three different microdomains: the matrix, intermembrane (IMS) and outer membrane space (OMS). Cells co-expressing one of these three targeted constructs with cytosolic mCherry were imaged before and after addition with a mild treatment of digitonin to permeabilize membranes along with a fluorescent quencher, trypan blue, to confirm mitochondrial microdomain localization^33^ (**Fig. 2A**). The fluorescence of microdomain-specific HyPer7 was compared to the reference fluorescent quenching of mCherry after over 15 min (1000 sec), when microdomain-specific differences in quenching rates would be apparent. As expected, quenching was dependent on targeting location, whereby HyPer7 was quenched slowest in the matrix and fastest in the OMS, proportional to the number of membrane layers enclosing that compartment (**Fig. 2B**).

**Figure 2:**
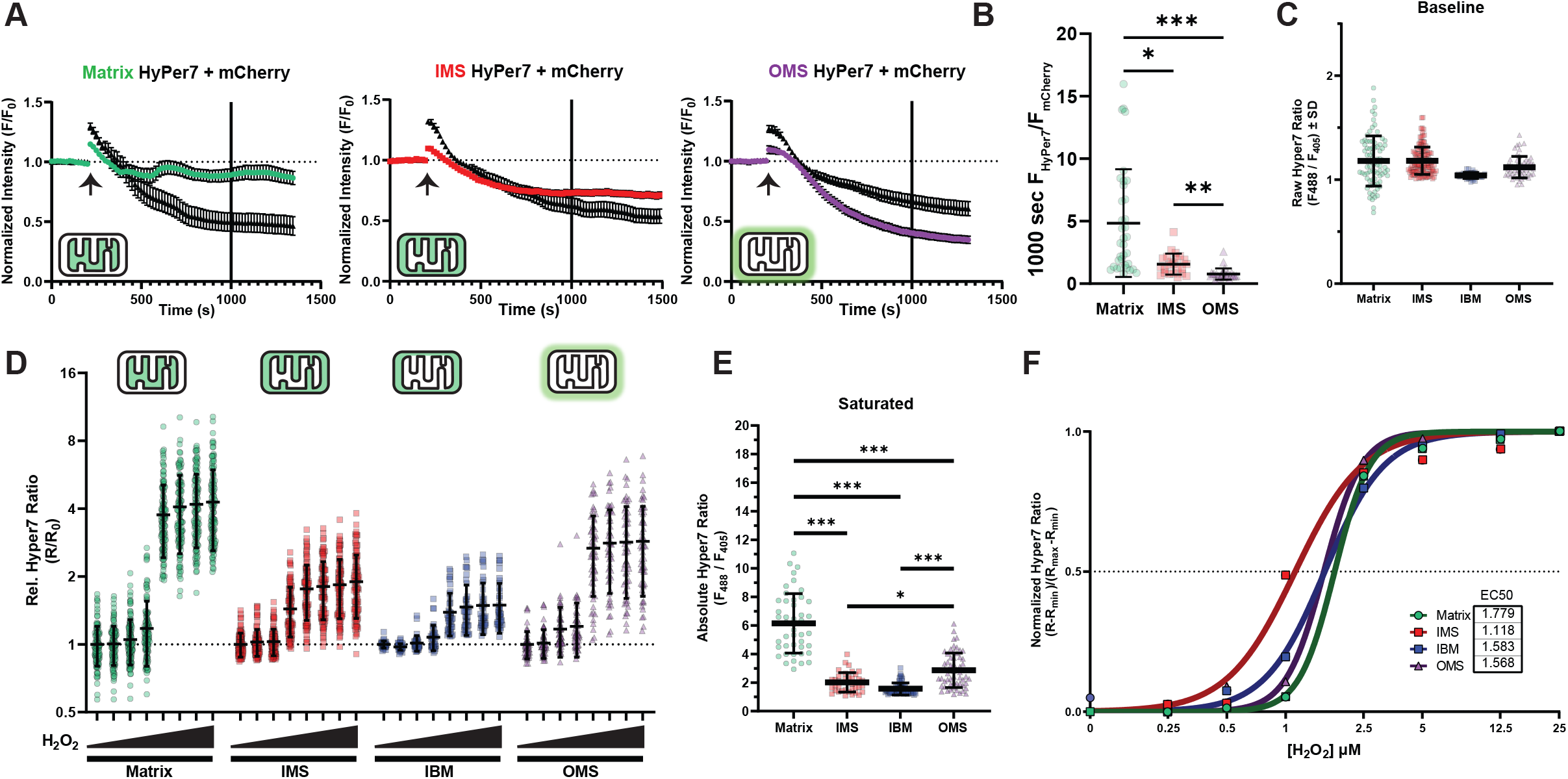
Mitochondrial microdomain-specific responses to exogenous hydrogen peroxide. A. Normalized microdomain-targeted HyPer7 and mCherry signals per cell relatively to baseline prior to treatment with digitonin and trypan blue. N > 30 cells per group. B. Ratio of microdomain-targeted HyPer7 to mCherry per cell. One-way Kruskal-Wallis test with Dunn’s *post-hoc* multiple comparisons correction. C. Absolute HyPer7 ratio per microdomain at baseline. N = 46 – 125 cells. D. Relative HyPer7 ratio response to increasing concentrations of hydrogen peroxide from 0.25 to 25 μM normalized to baseline per microdomain. N = 46 – 125 cells, mean ± SD. E. Absolute HyPer7 ratio per microdomain at saturating H_2_O_2_ exposure. One-way Kruskal-Wallis ANOVA with Tukey post-hoc correction, N = 38 – 71 cells, mean ± SD. F.Nonlinear sigmoidal fit of relative microdomain-specific HyPer7 responses to H_2_O_2_ after baseline and saturation normalization. *p < 0.05, **p < 0.01, ***p < 0.001.

We validated this targeting approach by expressing microdomain-specific HyPer7 into a previously confirmed common background of *C. elegans* expressing mCherry targeted to the inter-boundary membrane (IBM) of mitochondria^30,34,35^ (**Extended Fig. 2A**). The mitochondrial IBM represents the IMS, but without cristae^36,37^. Fluorescent proteins targeted to the IBM present as hollowed mitochondria when imaged at high resolution, thereby offering a visual reference standard to test microdomain-specific expression. Fluorescent line-scans of single mitochondria expressing microdomain-specific HyPer7 and IBM-mCherry further supported proper localization into the matrix, IMS, and OMS compartments based on the distance between compartment-specific HyPer7 and IBM-mCherry (**Extended Fig. 2B-C**).

With evidence of appropriate microdomain-specific targeting, we proceeded to identify how each compartment responded to exogenous treatments of H_2_O_2_. Whereas baseline steady-state levels of HyPer7 oxidation were largely unchanged between compartments (**Fig. 2C**), exogenous treatment of increasing H_2_O_2_ strongly affected HyPer7 responses (**Fig. 2D**). HyPer7 intensity at saturating levels of H_2_O_2_ significantly differed between microdomains (**Fig. 2E**). When normalized to microdomain-specific maxima, HyPer7 similarly responded to exogenous H_2_O_2_ treatment where the EC_50_ in each compartment ranged between 1 to 2 μM H_2_O_2_ with sharp saturation beyond 2.5 μM (**Fig. 2F**). Given the similarity between the normalized H_2_O_2_-response curves between microdomains, interpolating the exogenous H_2_O_2_ concentration that induced a given HyPer7 intensity change per microdomain seemed reasonable.

HyPer7 is reportedly pH-resistant^28^, but pH between mitochondrial microdomains can range up to a single unit^35^. To test whether the change in maximal HyPer7 response was dependent on microdomain-specific pH differences, we mutated the ROS-sensitive Cys121 residue to control for fluorescent changes independent of oxidation^28^. Baseline HyPer7(C121S) intensity further confirmed microdomain-specific targeting, as matrix-HyPer7(C121S) properly indicated a higher pH compared to IMS-HyPer7(C121S) (**Extended Fig. 3A**). Compared to HyPer7, the HyPer7(C121S) variant exhibited a dampened response in both the matrix and IMS to treatment with 100 μM H_2_O_2_ and a heightened response to 40mM NH_4_Cl to raise mitochondrial pH^38,39^ (**Extended Fig. 3B-C**). Treatment with saturating levels of H_2_O_2_ after NH_4_Cl incubation still led to an increase in HyPer7 signal comparable to purely H_2_O_2_-treated cells (**Extended Fig. 3D**), indicating pH has a minimal effect on HyPer7 ROS responses. Overall, the different responses of HyPer7 biosensor activity between mitochondrial compartments was not explained by differences in pH.

**Figure 3:**
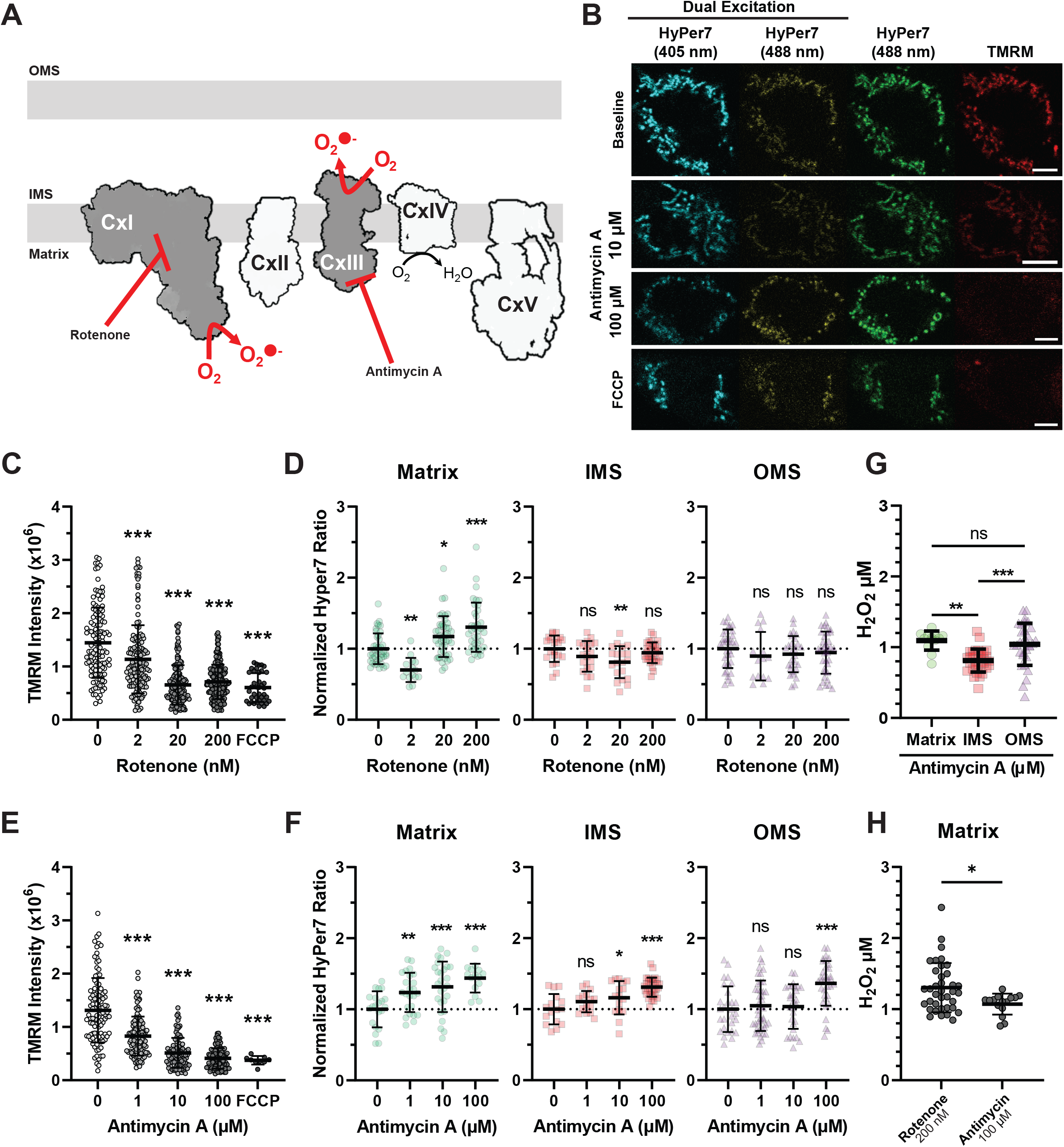
Microdomain-specific ROS diffusion to canonical mitochondrial toxins. A. Schematic of mitochondrial ETC toxins producing ROS at distinct sites. B. Representative images of dually excited HyPer7 and pseudo-simultaneous imaging of HyPer7 488 and TMRM in response to drug treatments. Scalebar denotes 5 μm. C. TMRM responses of cells treated with increasing concentrations of rotenone and FCCP. One-way Kruskal-Wallis ANOVA with Tukey post-hoc correction, N = 25 – 59 cells, mean ± SD. D. Relative HyPer7 ratio responses to ROS produced at complex I through increasing rotenone concentrations. One-way Kruskal-Wallis ANOVA with Tukey post-hoc correction, N = 15 – 59 cells, mean ± SD. E. HyPer7 responses to rotenone normalized to microdomain-specific HyPer7 maximum signal. One-way Kruskal-Wallis ANOVA with Tukey post-hoc correction, N = 25 – 59 cells, mean ± SD. F.TMRM responses of cells treated with increasing concentrations of antimycin A and FCCP. One-way Kruskal-Wallis ANOVA with Tukey post-hoc correction, N = 15 – 45 cells, mean ± SD. G.Relative HyPer7 ratio responses to ROS produced at complex III through increasing antimycin A concentrations. One-way Kruskal-Wallis ANOVA with Tukey post-hoc correction, N = 15 – 40 cells, mean ± SD. H.Comparison of interpolated steady-state H_2_O_2_ levels between maximal concentrations rotenone and antimycin A in matrix-HyPer7 expressing cells. Two-tailed, Mann-Whitney test, mean ± SD, N = 15 – 37 cells. *p < 0.05, **p < 0.01, ***p < 0.001.

We hypothesized HyPer7 could reveal distinct ROS clearance rates between microdomains because ROS scavenging and redox cycling proteins differ in the environments^40–45^. Surprisingly, HyPer7 decay rates did not differ between compartments and even between most H_2_O_2_ concentrations, decaying at ≈0.003 HyPer7 R/R_0_ per sec (**Extended Fig. 4A-B**). Overall, these data show HyPer7 responds to low micromolar exogenously added H_2_O_2_, but a more endogenous system was needed to analyze ROS dynamics in a microdomain-specific context.

**Figure 4:**
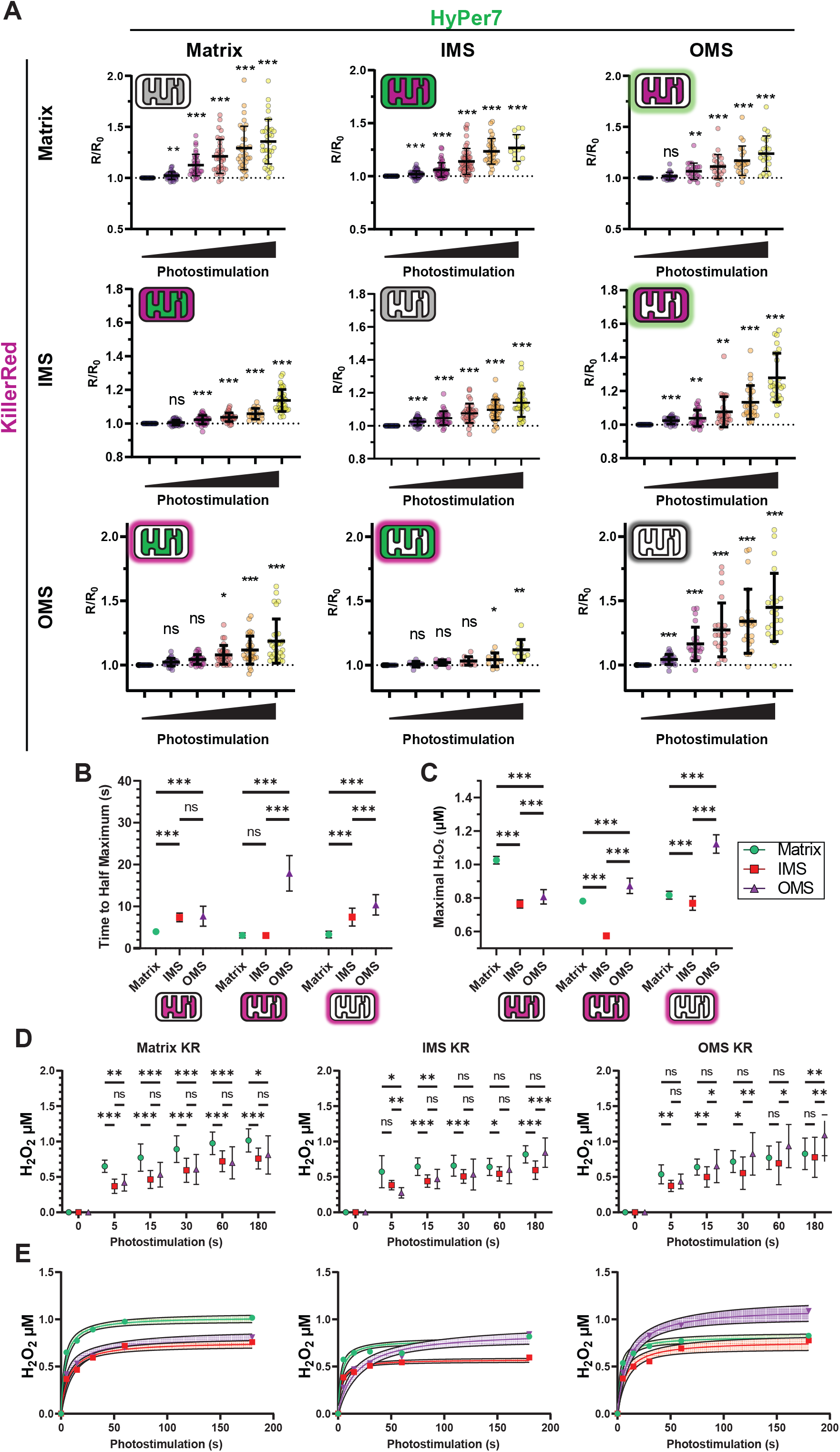
Diffusion kinetics of mitochondrial microdomain-specific ROS. A. HyPer7 normalized ratio responses with KillerRed co-expressed in all mitochondrial microdomains. KillerRed photostimulation ranged from 5 to 180 sec. One-way, repeated measure, mixed-effects ANOVA with Geisser-Greenhouse correction and Dunnett post-hoc correction, N = 21 - 55 cells, mean ± SD. B. Time to half maximum (T_1/2_) for microdomain-targeted HyPer7 and KR experiments. Two-way ANOVA with Tukey *post-hoc* correction, mean ± SD, N = 21 – 55 cells. C. Maximal interpolated steady-state H_2_O_2_ concentration of mitochondrial microdomains following microdomain-specific KR photogeneration of H_2_O_2_. Two-way ANOVA with Tukey *post-hoc* correction, mean ± SD, N = 21 – 55 cells. D. Interpolated apparent H_2_O_2_ concentrations in each microdomain from A. Two-way, repeated measure, mixed-effects ANOVA with Geisser-Greenhouse correction and Tukey *post-hoc* correction, N = 21 - 55 cells, mean ± SD. E. Interpolated apparent H_2_O_2_ concentrations as in B but depicted as nonlinear fit curves (all R^2^ > 0.6) across all photostimulation timepoints. *p < 0.05, **p < 0.01, ***p < 0.001.

### Measuring endogenously produced ROS diffusion between microdomains

Respiratory complexes along the ETC are involved in signaling metabolic state and regulating behavior^30^. Complexes of the ETC form microdomains of local ROS environments whereby oxidation of key cysteine residues can impair organismal responses to nutrient and oxygen deprivation which can mimic early stages of neurodegeneration^30,46,47^. Studying these key oxidative events is difficult, in part due to the lack of available tools to selectively oxidize mitochondrial microdomains with spatiotemporal resolution. One option to generate ROS in specific microdomains is by using mitochondrial toxins to block electron transport through the ETC at defined sites. For example, complex I inhibition by rotenone produces matrix ROS and inhibits ROS from reverse electron transfer, while complex III inhibition with antimycin A generates ROS in the inter-membrane space (IMS)^1,28,48–50^ (**Fig. 3A**).

HyPer7 has been previously reported to be sensitive enough to detect microdomain-specific ROS generation by respiratory chain toxins^12^. We treated cells expressing microdomain-targeted HyPer7 with increasing concentrations of rotenone and antimycin A to monitor ROS diffusion from these defined generation sites throughout microdomains. Dual-color excitation HyPer7 intensity levels were compared to the red fluorescent dye TMRM, an indicator of mitochondrial membrane potential, ΔΨ_m_. As rotenone and antimycin A block electron flow through the ETC, mitochondrial membrane potential drops as indicated by decreased TMRM signal. Simultaneously, superoxide is generated, ultimately producing H_2_O_2_ and oxidizing HyPer7 (**Fig. 3B**).

Simultaneous measurement of ΔΨ_m_ and ROS levels during acute ETC block revealed disparities in drug-mediated ROS diffusion through mitochondrial compartments. ΔΨ_m_ was highly sensitive to rotenone and antimycin A, as expected, where 2 nM and 1 μM, respectively, were enough to significantly lower ΔΨ_m_ (**Fig. 3C and F**). At higher concentrations, ΔΨ_m_ was fully impaired and comparable to full ΔΨ_m_ destabilization due to treatment with the protonophore FCCP. At our lowest concentration of 2 nM, rotenone significantly lowered matrix-HyPer7 oxidation compared to baseline (**Fig. 3D**), likely by inhibiting complex I and decreasing ETC-generated ROS, limiting ETC superoxide production^51,52^. Importantly, rotenone only oxidized matrix-HyPer7 and not IMS-or OMS-HyPer7 at any concentration (**Fig. 3E**). Under these experimental conditions, rotenone never induced ROS diffusion out of the matrix. Contrastingly, IMS-generated ROS through antimycin A diffused and oxidized both matrix- and OMS-HyPer7, but only once matrix-HyPer7 saturated at higher concentrations of antimycin A (**Fig. 3G**). This finding led us to estimate the amount of H_2_O_2_ generated in each compartment interpolated based on the microdomain-specific HyPer7 response to the H_2_O_2_ standard curve.

Interpolating based on this microdomain-specific standard curve revealed that ETC toxins produced unequal amounts of H_2_O_2_ per microdomain. Antimycin A produced less H_2_O_2_ in the IMS (0.81 ± 0.16 μM) than that of the matrix (1.09 ± 0.13 μM) or OMS (1.04 ± 0.29 μM) (**Fig. 3G**). The maximal concentration used for rotenone generated 1.30 ± 0.34 μM in the matrix, significantly higher compared to antimycin A (**Fig. 3H**). Overall, these results demonstrated directionally selective diffusion between microdomains depending on the local antioxidant state.

### Mapping the diffusion of endogenously generated H_2_O_2_ between mitochondrial microdomains

Respiratory toxins induce prolonged generation of ROS, but physiological ROS can be transient. To induce transient ROS and probe whether ROS can differentially diffuse between mitochondrial microdomains, we co-expressed HyPer7 and KR tagged to either the matrix, IMS, or OMS. These nine experimental conditions encompass ROS generation and diffusion between every mitochondrial microdomain. Cells co-expressing microdomain-specific HyPer7 and KR were continually imaged for nearly 20 min through increasing durations of full-frame photostimulation of green light at a fixed irradiance (0.42 mW/mm^2^, **Extended Fig. 4A-B**). Each combination of HyPer7 and KR microdomain targeting revealed light dose-dependent effects of KR photostimulation which saturated at longer durations of 180 sec (**Fig. 4A**). Longer photostimulation times exhibited weaker HyPer7 responses, likely because of oxidative damage to the KR protein which prevented further photosensitization^25,31,32^.

Interestingly, the rise times (time to half maximal HyPer7 response, T_1/2_), between compartments varied greatly (**Fig 4B**). Matrix KR led to strong HyPer7 responses in all microdomains, but the matrix (4.00 ± 0.68 sec) reached T_1/2_ faster than the IMS and OMS (7.38 ± 1.02 and 7.67 ± 2.38 sec, respectively). Consistent with our earlier results with antimycin A (**Fig. 3F-G**), ROS generated in the IMS with IMS-KR first led to a rapid increase in matrix-HyPer7 oxidation over IMS and OMS. The T_1/2_ rise time from IMS-KR was equal in the matrix (3.09 ± 0.61 sec) compared to the IMS (3.05 ± 0.45 sec), with a significantly delayed increase in the OMS (17.91 ± 4.24 sec). ROS generated at the OMS through photostimulation of OMS-KR more quickly diffused into the matrix (T_1/2_ risetime 3.30 ± 0.78 sec) over the IMS (7.46 ± 2.14 sec) and even OMS (10.37 ± 2.44 sec). These data suggest ROS made in the matrix diffuses into the IMS, which rapidly releases into the OMS. ROS made in the IMS initially diffuses into the matrix before a delay, ultimately being released out of the mitochondria into the OMS. ROS made in the OMS, in contrast, diffuses quickest into the matrix before a lag time where it eventually oxidizes the IMS. Overall, these data support the directionally selective diffusion of ROS within mitochondria microdomains.

To account for microdomain-specific response curves of HyPer7 and to enable accurate cross-compartment kinetic comparisons, we interpolated the concentrations of exogenous H_2_O_2_ which would approximate the HyPer7 response to KR photostimulation per microdomain (**Fig. 4C-E**). Matrix KR photostimulation culminated in a greater maximal steady-state of H_2_O_2_ concentration of 1.03 ± 0.02 μM in the matrix versus 0.76 ± 0.02 and 0.81 ± 0.04 μM in the IMS and OMS, respectively. As photostimulation duration increased, the OMS-HyPer7 response increased to supersede the matrix response by 180 sec of photostimulation. At this maximal photostimulation duration, ROS generated in the IMS led to the highest H_2_O_2_ concentration in the OMS (0.87 ± 0.05 μM) compared to the IMS (0.57 ± 0.01 μM) and matrix (0.78 ± 0.02 μM), though the matrix remained significantly higher than the IMS. At maximal photostimulation duration of OMS-KR, this pattern was reversed whereby steady-state H_2_O_2_ was highest in the OMS (1.12 ± 0.05 μM) relative to the matrix (0.82 ± 0.02 μM) and IMS (0.77 ± 0.04 μM), as originally expected. While this data elucidated initial directionality in diffusion between mitochondrial compartments, we further asked whether the decay kinetics of ROS in each compartment would reflect the ROS microenvironments present at each microdomain.

### All-optical recording of ROS release kinetics from the matrix in single mitochondria

Redox scavenging and signaling machinery differ in expression among mitochondrial microdomains and the cytosol^53–56^. HyPer7 oxidation is partly dependent on the scavenging mechanisms competing for H_2_O_2_, whereas the decay rate of HyPer7 fluorescence following oxidation depends on the local redox machinery available to re-reduce HyPer7^12,28^. Having found differences in the approximate steady-state H_2_O_2_ levels produced by KR stimulation in each microdomain, we tested whether the HyPer7-KR system would be sensitive enough to detect differences in microdomain-specific rise and decay rates of H_2_O_2_ due to differing scavenging mechanisms. To accomplish this, we repeatedly induced ROS production in the matrix of single mitochondria through matrix-KR photostimulation and compared recordings of HyPer7 oxidation present in different mitochondrial microdomains. Given the technical limitations of slow light path switching between dual-color excitation of HyPer7 and single-color KR imaging, we resorted to simultaneous single-color excitation of HyPer7 (488 nm) and KR (561 nm) every 5 sec for over 20 minutes of recording followed by offline normalization of each individual mitochondrion’s HyPer7 and KR intensity to its own area per frame to mitigate three-dimensional focal drift over time as in **Fig. 1**.

Broad details of ROS release from the matrix in mitochondria emerged when the oxidation of each microdomain-targeted HyPer7 were analyzed as fluorescent change over baseline. Spatially restricted ROS production through photostimulation of matrix-KR was confirmed by comparing the responses of mitochondrial populations expressing microdomain-targeted HyPer7 as a function of distance from a single pulse of 561 nm at a fixed point (**Fig. 5A**). As the distance between mitochondria and the fixed stimulation point increased, the resulting photobleaching of KR and subsequent oxidation of HyPer7 decreased. Matrix-HyPer7 failed to respond to matrix-KR photostimulation at approximately 12 μm, whereas IMS- and OMS-HyPer7 failed to respond at roughly 6 μm and 1 μm, respectively. H_2_O_2_ generated by matrix-KR photostimulation produced decreasing HyPer7 responses as the targeted microdomains were further from the matrix, indicating the single pulse of ROS filtered through continuing antioxidant machinery per microdomain.

**Figure 5:**
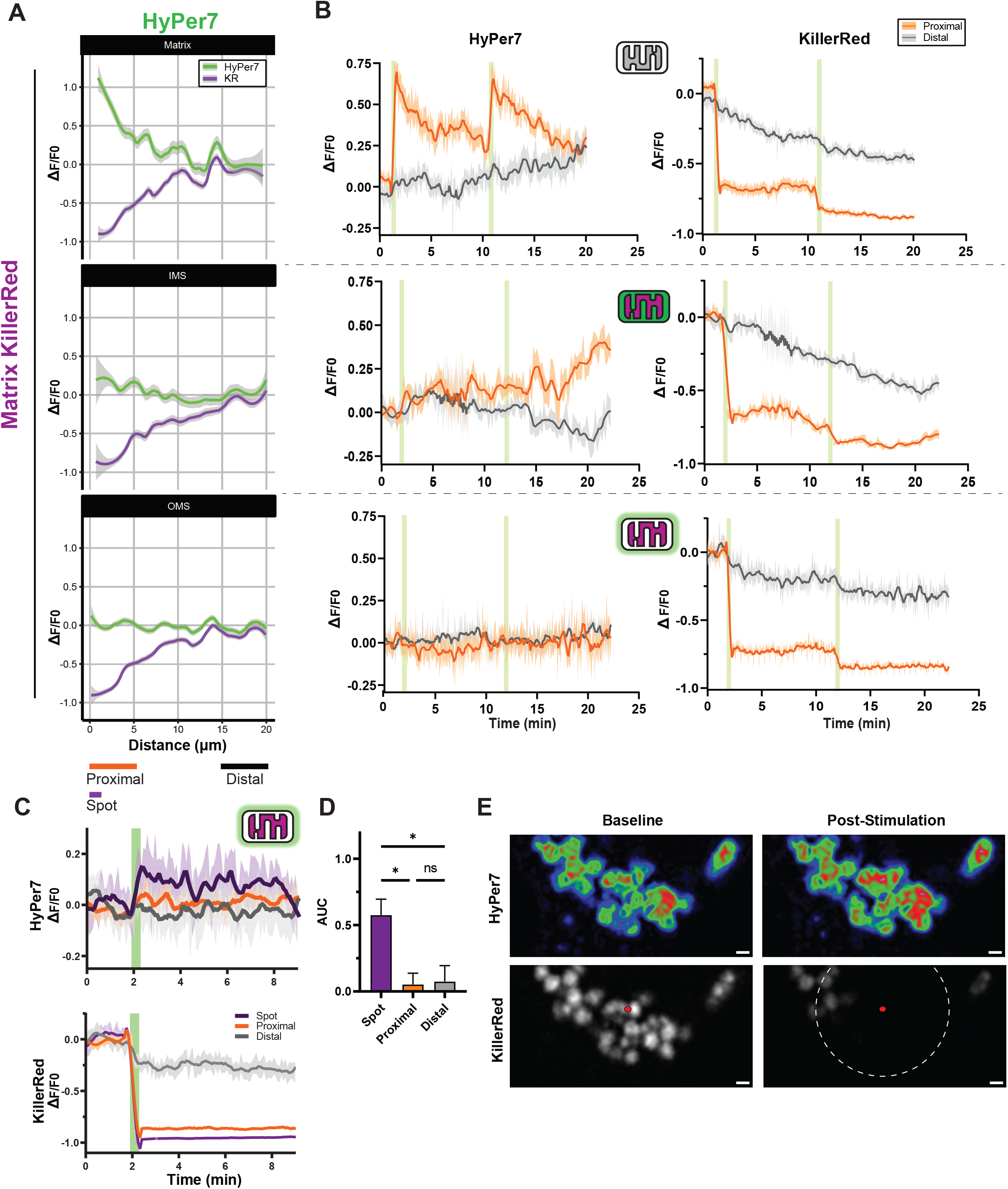
Dynamics of single mitochondrial matrix ROS release. A. Fluorescent change over baseline of single mitochondria in cells co-expressing various microdomain-targeted HyPer7 with matrix-KillerRed. Data taken from 12 frames (one minute) following the first 5 sec photostimulation pulse and displayed as a function of distance from the spot stimulation point. Data expressed as mean ± 95% CI. N > 60 individual mitochondria per frame, on average. B. Average traces of fluorescent change over baseline of single mitochondria co-expressing microdomain-targeted HyPer7 and matrix-KillerRed following repeated photostimulation. Mitochondrial populations were separated based on distance to spot stimulation, where proximal and distal roughly approximates under 5 μm and over 15 μm, respectively. Data expressed as smoothed mean ± SEM. N > 60 individual mitochondria per frame, on average. C. Fluorescent change over baseline of single mitochondria expressing OMS-HyPer7 and matrix-KillerRed following a single, longer duration (30 sec) photostimulation pulse of KillerRed. Mitochondrial populations were separated as proximal and distal, but also included a “spot” subpopulation of mitochondria within 1.5 μm of the stimulation point. Data expressed as smoothed mean ± SEM, N = 6, 27, and 52 individual mitochondria for spot, distal, and proximal mitochondrial populations per frame, on average. D. Area under the curve analysis of C. Data expressed as mean ± SEM, one-way ANOVA with Tukey post-hoc correction, N = 4 independent experiments. E. Representative images of mitochondria in a cell expressing OMS-HyPer7 (rainbow LUT applied) and matrix-KillerRed (grayscale) before and after photostimulation. Spot stimulation point depicted as a red circle, 5 μm surrounding area as a dashed white circle. Scalebar denotes 1 μm. *p < 0.05, **p < 0.01, ***p < 0.001. Note: Matrix-HyPer7 data in b) taken from Figure 1 and re-expressed as dF/F.

Contrary to whole-cell KR photostimulation (**Fig. 4**), spatially restricting acute photostimulation of KR revealed the variation in microdomain-specific responses over time. Matrix-HyPer7 exhibited repeatable stimulation, but the IMS-HyPer7 response was initially dampened until nearly five minutes after the second light pulse (**Fig. 5B**). We reasoned that this delayed increase in oxidation resulted from the saturation of IMS-specific antioxidants such as cytochrome c and the relatively lower amounts of available redox machinery at the IMS, resulting in ROS-induced ROS release (RIRR) in the IMS which was not apparent in the matrix^56–58^. Notably, this delayed increase in IMS HyPer7 oxidation coincided with the second pulse decay of matrix HyPer7 and increase in mitochondrial motility. Interestingly, neither pulse of matrix KR photostimulation led to increased OMS-HyPer7 responses, indicating this acute, spatially restricted ROS generation paradigm did not overwhelm antioxidant machinery enough to oxidize OMS-HyPer7 and measurably alter its fluorescence.

To test whether we could detect release of matrix ROS into the OMS in single mitochondria, we increased the duration of photostimulation pulse from 5 sec to 30 sec. Increasing the photostimulation time and further subdividing mitochondria proximal to the spot stimulation point to within 1.5 μm of the spot stimulation point (denoted as “spot,” compared to proximal of less than 5 μm and distal as over 15 μm away) revealed a marked increase in OMS-HyPer7 oxidation that remained for nearly 10 min (**Fig. 5C**). Area under the curve (AUC) analysis revealed this prolonged stimulus oxidized OMS-HyPer7 in a highly spatially restricted manner (**Fig. 5D-E**). These data suggest that coupling expression of microdomain-targeted HyPer7 and KR with single mitochondrial photostimulation can reveal the spatiotemporal diffusion kinetics of endogenously produced H_2_O_2_ between compartments and measure the release of matrix ROS into the cytosol without other alterations.

## Discussion

### Limitations in all-optical ROS control

Here, we use microdomain-targeted expression of the H_2_O_2_ biosensor HyPer7 and ROS photogenerator KillerRed to measure the kinetics of ROS generation and diffusion throughout mitochondrial microdomains. One limitation of this study is the inability to directly control for microdomain-specific differences in volume or microdomain insertion of targeted proteins. Given the apparent differences in microdomain-specific HyPer7 responses, controlling for these differences was crucial for inter-microdomain comparison. To account for this, all intensity measurements described here involve multiple measures of intra-experimental normalization. For example, HyPer7 ratiometric intensities were used in most comparisons which mitigated differences in biosensor expression and insertion. During single mitochondrial imaging experiments, single excitation HyPer7 intensities per mitochondrion were normalized to that same mitochondrion’s area in each frame, which similarly limited expression-based confounds. Cross-compartment, all-optical perturbation of ROS using HyPer7 and KR were normalized first to each individual cell’s KR expression, and then by measuring ratiometric HyPer7 intensities relative to the baseline of the same cell following photostimulation. Moreover, interpolated H_2_O_2_ concentrations was based on microdomain-specific responses, further controlling for any apparent differences. Together with multiple factor assessment of microdomain-specific targeting and HyPer7 pH dependence, these methods attempted to account for inherent bias in technical approaches. It remains possible, though unlikely, that fundamental microdomain-specific changes in biosensor responses bias our results despite these normalization approaches. Thus, interpretation of the resulting H_2_O_2_ concentration we report herein should be carefully considered given the technical and biological complexities of measuring H_2_O_2_ dynamics *in situ*^1,59^.

### Estimating steady-state H_2_O_2_ levels in live cells

We use microdomain-specific standard curves of exogenously added H_2_O_2_ to compare HyPer7 responses due to toxins or KR photogeneration of ROS between microdomains. Since H_2_O_2_ diffuses through membranes^17,20,60,61^ and local redox scavengers such as the peroxiredoxins shift HyPer7 responses by competing for H_2_O_2_ ^12,28^, we consider these microdomain-specific responses a reflection of apparent redox-state in that compartment. In other words, an increase in HyPer7 signal means the antioxidative machinery in that compartment could no longer prevent ROS from oxidizing proteins such as HyPer7, reducing any oxidized thiols, and/or dismutase activity in that compartment facilitated H_2_O_2_ production. Moreover, antioxidant buffering systems such as the peroxiredoxin and thioredoxin systems responding to H_2_O_2_ would continually diffuse and equilibrate between cellular compartments during ROS clearance^12^. Therefore, a lack of HyPer7 oxidation does not necessarily indicate the lack of ROS diffusion^62,63^. This is best exemplified in the IMS, where the relatively lower concentration of SOD compared to the matrix and cytosol, but with an increase in cytochrome c antioxidant levels, is predicted to sequester superoxide prior to dismutation into H_2_O_2_ ^40,56,64,65^. These differences could ultimately limit H_2_O_2_ oxidation of HyPer7 and explain the consistently lower apparent H_2_O_2_ levels detected by HyPer7 in the IMS in our experiments.

Moreover, since the concentration of exogenously added H_2_O_2_ is likely an over-estimate of intracellular and subcellular accumulation, interpolating local concentrations of H_2_O_2_ based on HyPer7 measurements can only provide an upper-bound estimate. Other studies have modeled a 390 to 650-fold difference between extracellular and intracellular H_2_O_2_^62,63^ and calculated a basal steady-state of 2 to 4 nM in mitochondria^9^. Our experiments never produce a steady-state change approximating greater than 2 μM H_2_O_2_ treatment. This suggests the apparent mitochondrial accumulation of H_2_O_2_ corresponded to between a 3 to 5 nM increase above this baseline, though this number is likely specific to both cell type and metabolic conditions. Acute ROS generation experiments using toxins and KR reported here never caused overt cell death during imaging and had only begun to overload redox machinery between compartments. Therefore, we predict our estimated 3 to 5 nM increase in mitochondrial H_2_O_2_ delineates the transition from oxidative eustress toward distress. Higher levels of ROS production induce mitochondrial aggregation, dysfunction, mitophagy and ultimately cell death^25,31,34,68–71^. Future studies in alternative models, scavenging, and metabolic or disease conditions may further investigate the responses to changes in ROS.

### Intracellular ROS spreads through the mitochondrial network on the minute timescale

Previous reports have indicated ROS-mediated hyperfusion acts to prevent pathological accumulations of ROS in the mitochondrial network^72–75^. Here, we show that spatiotemporally limited pulsing of matrix ROS generation induced transient mitochondrial hyperfusion and intracellular mixing of ROS through the active movement of mitochondria. The first pulse of matrix ROS induced local hyperfusion. The second pulse induced less transient hyperfusion, but also increased ROS levels found in distal mitochondria relatively far from the site of ROS photogeneration. This network ROS spread was delayed by several minutes following the second pulse of ROS itself and was limited by cell-type. However, in human cells, this inter-network spread occurred during the decay of matrix HyPer7 and during the increase in IMS HyPer7 oxidation, indicating ROS was being released from the matrix. We speculate this inter-mitochondrial spread of ROS acts to dilute ROS contents over a greater population of mitochondria, both decreasing the damage of any singular mitochondrion but also hastening the global cell response to oxidative distress. An important extension of this work would be investigating mitochondrial ROS spread in cells with unique morphology, such as neurons, where mitochondria can be trafficked over relatively large distances.

Whether this increase in HyPer7 oxidation in distal mitochondria is produced by the inter-mitochondrial exchange of H_2_O_2_ or oxidized HyPer7 is unknown, but either possibility supports inter-mitochondrial content exchange to ostensibly lower any single mitochondrion’s damage due to ROS. Given that the spatial positioning of mitochondria influence subcellular redox status^76^, this transient increase in fusion followed by mitochondrial motility could work to prevent aberrant mtDNA and protein oxidation in response to mild ROS stimulus^73,77^. This ROS-mediated pulse of mitochondrial motility and spread may contribute to region-specific differences in oxidative vulnerability, such as in neurons^68,78^.

### Directional ROS flow between mitochondrial compartments prior to release into the cytosol

Many studies have used rotenone and antimycin A to generate and accumulate ROS to study downstream physiological and pathological effects. Our acute treatments of rotenone and antimycin A across low to high doses generated steady-state HyPer7 oxidation levels which approximated treatments of under 2 μM H_2_O_2_. Notably, care should be taken not to equate treatment of 10 or more μM H_2_O_2_ with short-term treatment of rotenone or antimycin A given these differences. Our lowest dosage of rotenone (2 nM) lowered matrix-HyPer7 oxidation, consistent with the theory that low dose complex I inhibition decreases ROS and provides physiological benefits that extend lifespan^79^. Across all dosages, however, rotenone did not lead to oxidized HyPer7 in the IMS and OMS. As said previously, this lack of HyPer7 oxidation does not necessarily indicate the lack of ROS spread, but instead that antioxidant systems in these compartments adequately cleared any diffusing ROS and prevented the oxidation of HyPer7. In contrast, generation of ROS in the matrix through matrix-KR photostimulation led to equal increase in HyPer7 oxidation in the IMS and OMS. This generally agrees with the “flood-gate” model of ROS dynamics^62,63^, where a prolonged ROS stimulus (e.g., rotenone) can be cleared over time, but a sudden larger insult of ROS (e.g., pulsed KR stimulation) overwhelms redox machinery and leads to diffusion between compartments and ROS release into the cytosol.

Unlike matrix ROS, IMS ROS diffuses in a directionally selective manner depending on concentration. Increasing dosages of antimycin A induced increasing HyPer7 oxidation in the matrix before being released from mitochondria and diffusing into the OMS. Using KR as an optically-controlled photosensitizer allowed finer temporal control of this directional selectivity, where the lowest photostimulation pulse induced mostly inward ROS diffusion to the matrix. Lengthening photostimulation duration led to saturation of matrix HyPer7 and subsequent OMS-HyPer7 oxidation. It remains to be seen if a biosensor can detect basal differences in ROS between mitochondrial compartments as predicted based on physiological site-specific production of ROS along the ETC.

Considerable debate has focused on whether H_2_O_2_ generated within the mitochondrial matrix can be released into and impact other cellular compartments^28^. Here, we maintained constant metabolic conditions of relatively low, near-physiological 3-5 mM glucose, compared to the more common 10-25 mM concentrations seen in culture media^12^ which protected mitochondrial morphology and limited downstream effects of hyperglycemia such as ROS accumulation during imaging experiments^80–83^. Using systematic, all-optical cross-compartment detection and generation of H_2_O_2_, we uncover the spatiotemporal release dynamics of matrix-generated ROS into the cytosol without metabolic or redox machinery perturbation. As our experiments with OMS-KR indicate, ROS generated on the outer membrane of mitochondria is rapidly internalized in the matrix and can oxidize HyPer7, even during the shortest photostimulation pulse of KR. Our single mitochondrial photostimulation experiments using matrix-KR are consistent with this idea, where strong pulses of matrix ROS can be released and diffuse between mitochondria across roughly 1-2 μm to oxidize matrix-HyPer7. This is consistent with the estimated lifetime diffusion of H_2_O_2_ in the cytosol of roughly 1 ms^84,85^.

Together, these data (summarized in **Extended Figure 6**) reveal the diffusion kinetic and directionality details of endogenously-produced ROS between mitochondrial compartments in living mammalian cells. Beyond these parameters, we suggest a model whereby ROS generation causes prolonged effects on mitochondrial morphology and ROS steady-state levels due to local ROS release from mitochondria into the cytosol, where mitochondrial hyperfusion and motility lead to cell-wide changes in redox state. These findings further support how understanding ROS dynamics between mitochondrial microdomains can reveal how ROS can act as a metabolic signal which propagates within the mitochondrial network, influencing cell-wide physiological responses over short timescales.

## Methods

### Plasmid Construction

Matrix- and IMS-HyPer7 was originally received as a gift from Vsevolod V. Belousov (Addgene #136470 and 136469). KR from Evrogen (#FP963, pArrestRed) was subcloned in another study^34^. To alter tags and vectors for use in cells and worms in this study, constructed were subcloned using in vivo assembly (IVA)^86^. HyPer7(C121S) was mutated using point mutagenesis. All primers used are supplied in **Supplementary Table 1**. Microdomain targeting was accomplished using N-terminal fusions of tags to the matrix using 2xCOX8A (tag corresponds to Cox8A N-terminal residues 1-25, from Addgene 136470), the IMS using SMAC (residues 1-59, from Addgene 136469), the IBM using IMMT (residues 1-187^34^), and the OMS using TOMM20 (residues 1-55, from Addgene 66753).

### Cell culture and transfection

HEK293T cells (System Biosciences) and mouse embryonic fibroblasts (MEFs, ATCC) and were grown at 37°C with 5% CO_2_ in DMEM medium (glucose 4.5 g/L) supplemented with 10% fetal bovine serum (FBS), 2 mM GlutaMax (ThermoFisher), and penicillin/streptomycin and plated onto MatTek 35 mm glass-bottomed dishes (20 mm, No. 1.5) at approximately 200,000 cells ml^-1^ two days before imaging and one day before transfection. For TMRM and drug experiments, 50,000 cells ml^-1^ were plated into 8-well chambered No. 1.5 coverslips (Ibidi). DNA transfections for HEK cells used PolyJet (SignaGen Laboratories), whereas MEF cells used TransIT-X (Mirus) following manufacturer protocol with varied DNA concentrations. For both cell types, each plate was transfected using 0.15 μg of each microdomain-targeted construct and brought to 1 μg with empty pcDNA3.1+ vector.

### Cell imaging

Transfected cells had media replaced with prewarmed HBSS imaging buffer composed of (in mM): 5.33 KCl, 0.44 KH_2_PO_4_, 4.16 NaHCO_3_, 137.9 NaCl, 0.33 Na_2_HPO_4_ supplemented with 20 HEPES and 3-5 glucose for at least 20 minutes prior to imaging. For some experiments, TMRM (25 nM) was added to imaging media. For respiratory toxin experiments, varying concentrations of rotenone (Sigma) and antimycin A (Sigma) were included in imaging buffer. Cells treated with rotenone and antimycin A were incubated at 37°C with 5% CO_2_ for 30 and 60 min, respectively, prior to imaging.

Cells were imaged using an inverted Nikon A1R HD microscope with 405, 488, 561 nm laser lines with a CFI Apochromat TIRF 60x oil objective. HyPer7 was imaged in dual excitation, single emission setup with 405 and 488 nm excitation unless otherwise stated. KillerRed was imaged using the lowest possible 561 nm excitation. Images taken using resonant scanning used at least 2x line averaging which was kept constant for each experiment. Galvano scanning was used for single mitochondrial imaging with 11.67x optical zoom (19.3 px / μm). To ensure minimal drift, samples were allowed to equilibrate on the microscope for 5-10 min prior to imaging onset.

### KillerRed activation

Activation of KillerRed in the entire field of view used a white light Sola light engine filtered through a TexasRed filter (30.5%, 0.42 mW/mm^2^). Spot stimulation of KillerRed was targeted with a 561 nm laser (30% power, 88 μW, 30 Hz, 100% duty cycle for 5 sec stimulation).

### Mitochondrial topology

Cells expressing various microdomain-targeted HyPer7 constructs were co-transfected with cytosolic mCherry for use as a comparison standard between quenching experiments. HyPer7 and mCherry were imaged for 3 min baseline in imaging buffer, followed by two washes and replacement of media with prewarmed intracellular buffer (in mM): 10 NaCl, 130 KCl, 1 MgCl_2_, 1 KH_2_PO_4_, 2 succinic acid, and 20 HEPES (pH 7.4) supplemented with 0.5 mg/ml trypan blue and 20 μM digitonin. Experiments with movement artifacts due to media replacement were discarded so single cell traces of HyPer7 and mCherry signal could be obtained and compared to same-cell baseline throughout the entire recording. Cells which did not respond to digitonin were not analyzed.

### HyPer7 responses

Hydrogen peroxide was diluted with imaging buffer and kept on ice. Baseline measurements of HyPer7 steady-state in each microdomain was measured following baseline recording where cold imaging buffer was added to mock H_2_O_2_ treatment. Increasing concentrations of H_2_O_2_ were applied to cells and imaged for at least 3 min before treatment with the next concentration. For decay kinetic experiments, HyPer7 responses to some concentrations were imaged for 10-20 min prior to next addition of H_2_O_2_. Saturated responses of microdomain-targeted HyPer7 were taken from 100 μM additions of H_2_O_2_ either independently or at the end of the increasing H_2_O_2_ concentration curves. H_2_O_2_ concentrations were interpolated from averages of microdomain-specific responses of HyPer7 based on a sigmoidal curve where R^2^ > 0.99 for all curves.

### Single mitochondrial photostimulation

Single cells brightly expressing microdomain-targeted HyPer7 and KillerRed were chosen for spot stimulation and single mitochondrial analysis. A pre-determined spot in the cell was targeted for repeated stimulation and remained unmoved during imaging. The XY coordinate of the spot was marked for later analysis. Following 1-3 min baseline recording, two iterations of a 5 sec spot stimulation followed by 10 min recording were imaged. Since imaging KillerRed also leads to slight ROS photogeneration, the lowest possible 561 nm laser power (0.27% power, 2.2 μW) was used which still enabled single-mitochondrial resolution. No images underwent photobleaching or drift correction. Average traces of single mitochondria data were smoothed for graphical representation, but statistical analysis was done on raw data.

### Image analysis

Images were loaded into FIJI/ImageJ (v1.5.3) and background subtracted (radius = 50). Rarely, minor lateral drift was corrected for using the “correct 3D drift” plugin during long-term imaging. Cells which drifted substantially were discarded, but measurements prior to drift were included in analysis. Each cell was manually traced for region-of-interest (ROI) measurement of fluorescent intensities. Unless otherwise stated, the ratio of HyPer7 emission from 488 nm excitation was divided by the emission from 405 nm excitation and was normalized to same-cell baseline through H_2_O_2_ or photostimulation experiments. For microdomain H_2_O_2_ diffusion experiments, KillerRed intensity was obtained prior to photostimulation to normalize the same cell HyPer7 response to KillerRed expression. Cells with mitochondrial aggregation or other gross morphological deformities were not included in analysis.

### Mitochondrial morphology

Following background subtraction, images were smoothed with a gaussian filter (sigma = 2). Mitochondria were automatically segmented by a Weka segmentation classifier (v3.3)^87^ trained on manually drawn ROIs. TrackMate (v7)^88^ was used to report the sum of HyPer7 and KillerRed intensity, area, perimeter, and the major and minor elliptical axis dimensions for each mitochondrion for every frame. The Euclidean distance from the centroid of each mitochondrion per frame to the XY coordinate of spot stimulation was calculated. Data were then analyzed using custom code in R, where the HyPer7 and KillerRed intensities per mitochondrion were normalized to its area to limit artifacts based on Z-drift and protein expression differences. Form factor [perimeter^2^/(4π × area)] and aspect ratio (major / minor elliptical axis) were calculated for each mitochondrion in each frame. Rather than use an arbitrary distance cutoff since cellular size and shape, mitochondria were classified as proximal and distal based on that mitochondrion’s proportion to the maximum distance found of any single mitochondrion to the stimulation spot per cell. Proximal and distal mitochondria were defined as less than 30% and more than 70% of the maximum distance of any mitochondrion to the spot stimulation, respectively, which corresponded to approximately 5 and 15 μm. In MEFs, these values were changed to under 20% and over 50%, respectively. For statistical analysis of morphology, only timepoints in which proximal mitochondria were significantly different from both non-stimulated control and distal subpopulation mitochondria were reported.

### C. elegans imaging and single mitochondrial linescans

*C. elegans* were maintained at 20°C on plates containing nematode growth media (NGM) with OP50 bacterial as food as previously described^30^. Strains expressing microdomain-targeted HyPer7 with a common IMMT::mCherry background were generated in-house^34^. Staged L4 worms were grown on standard NGM plates and screened for bright expression of HyPer7. Prior to imaging, worms were placed onto a glass slide agar pad containing 10 mM tetramisole hydrochloride and sealed with a No. 1.5 glass coverslip 20 min prior to imaging. Worm hypodermal cell mitochondria were identified by morphology and imaged with 488 and 561 nm laser lines to visualize HyPer7 and mCherry, respectively. HyPer7/mCherry intensities across linescans manually drawn through individual mitochondria were analyzed in ImageJ.

### Statistics

All data were plotted and statistically analyzed in GraphPad Prism (v9) or R (v4.0) and first subjected to a normality test. Single variate analyses were run as either unpaired, two-tailed t-tests or Mann-Whitney U tests. Multivariate analyses were run as either parametric one-or two-way ANOVAs, nonparametric Kruskal-Wallis tests, or repeated measure, mixed-effect two-way ANOVAs with Geisser-Greenhouse correction. All multivariate tests used *post-hoc* corrections as described in the figure legend text. Outliers were excluded in TMRM response curves to rotenone and antimycin following a 1% ROUT test (9/420 cells) which had no effect on reported statistical comparisons. Statistical p values < 0.05 were considered significant with any p values under 0.001 truncated and reported as ***.

## Acknowledgements

APW laboratory is supported by grants from National Institutes of Health (R01 NS092558, R01 NS115906). Some strains were provided by the Caenorhabditis Genetics Center (CGC), which is funded by NIH Office of Research Infrastructure Programs (P40 OD010440). JE and GVWJ are supported by R01 AG073121. SAK current affiliation: Harvard University, Department of Neurobiology, Boston, MA 02115, United States of America.

## Author contributions

SAK and APW conceptualized and oversaw the project. SAK wrote the manuscript. SAK, NAS, LDLR, JH, MAF, AYW, AME, and JE performed experiments. GVWJ contributed resources and funding. All authors edited and approved the final manuscript.

## Data, Materials, and Code Availability

Data and plasmids generated in this study are available upon reasonable request. Custom R code used in this study will be released on Github following publication.

## Extended Figures

**Extended Figure 1.**
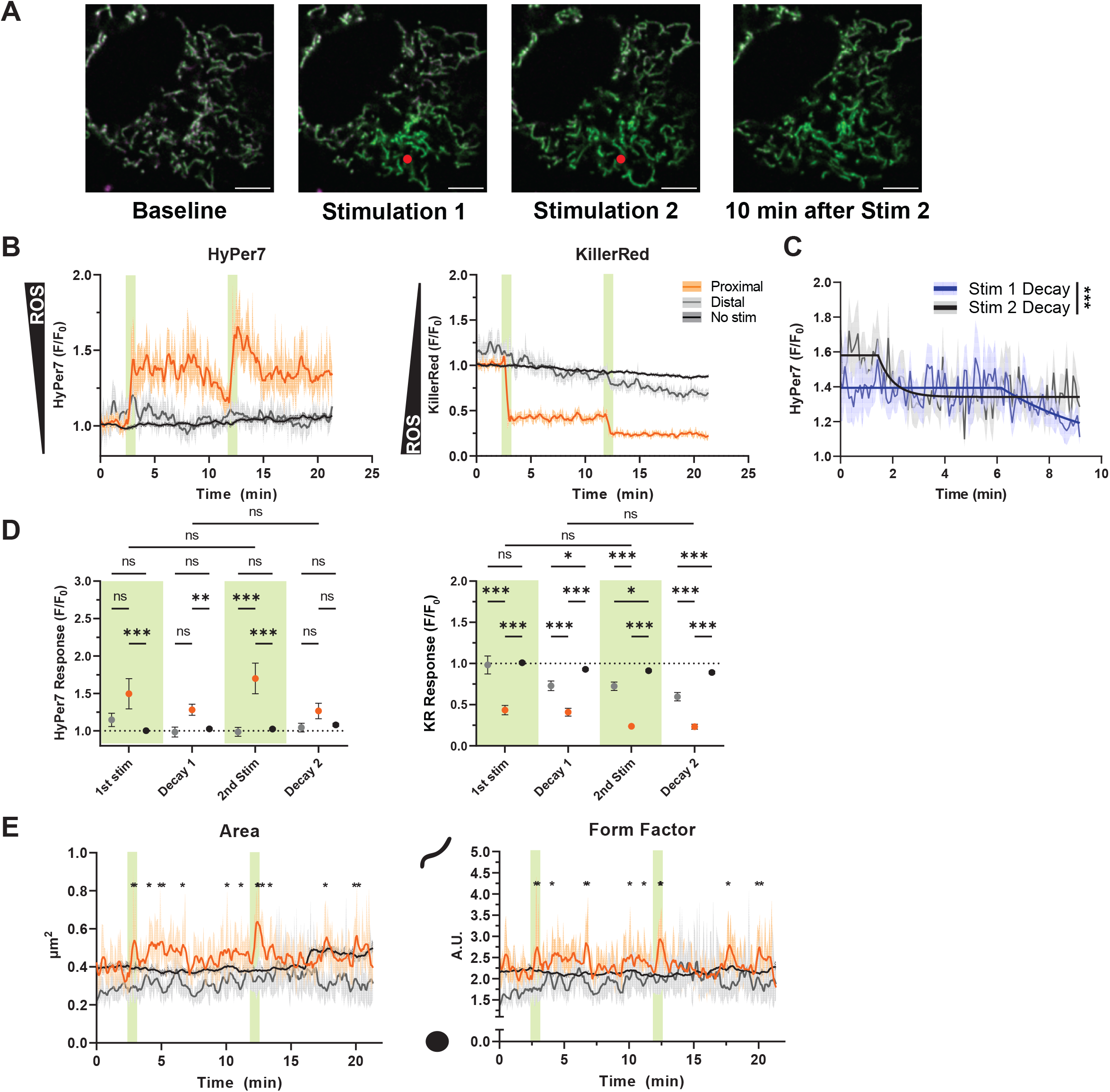
Spatiotemporal manipulation and monitoring of ROS in MEF single mitochondria. a) Representative image of Matrix-HyPer7 and Matrix-KR photostimulation in MEF cells. Scale bar represents 5 μm. b) Single mitochondrial responses of HyPer7 and KR intensities normalized to area and baseline, mean ± SEM, N = 20-334 mitochondria on average per group. c) Decay of stimulation 1 and 2 fitted with a nonlinear variable plateau followed by single phase decay, smoothed mean ± SEM. d) Comparison of HyPer7 and KR responses at first frame immediately following pulse 1, 10 minutes following first stimulation, immediately following pulse 2, and 10 minutes following second stimulation. Two-way ANOVA with Tukey *post-hoc* multiple comparisons, mean ± SEM. e) Measured mitochondrial morphological characteristics between subpopulations. Only significant timepoints where proximal measures were significantly different from both distal and no stimulation controls are shown. Two-way ANOVA with Tukey *post-hoc* multiple comparisons, mean ± SEM. Green bars indicate single frames of KR photostimulation. *p < 0.05, **p < 0.01, ***p < 0.001.

**Extended Figure 2.**
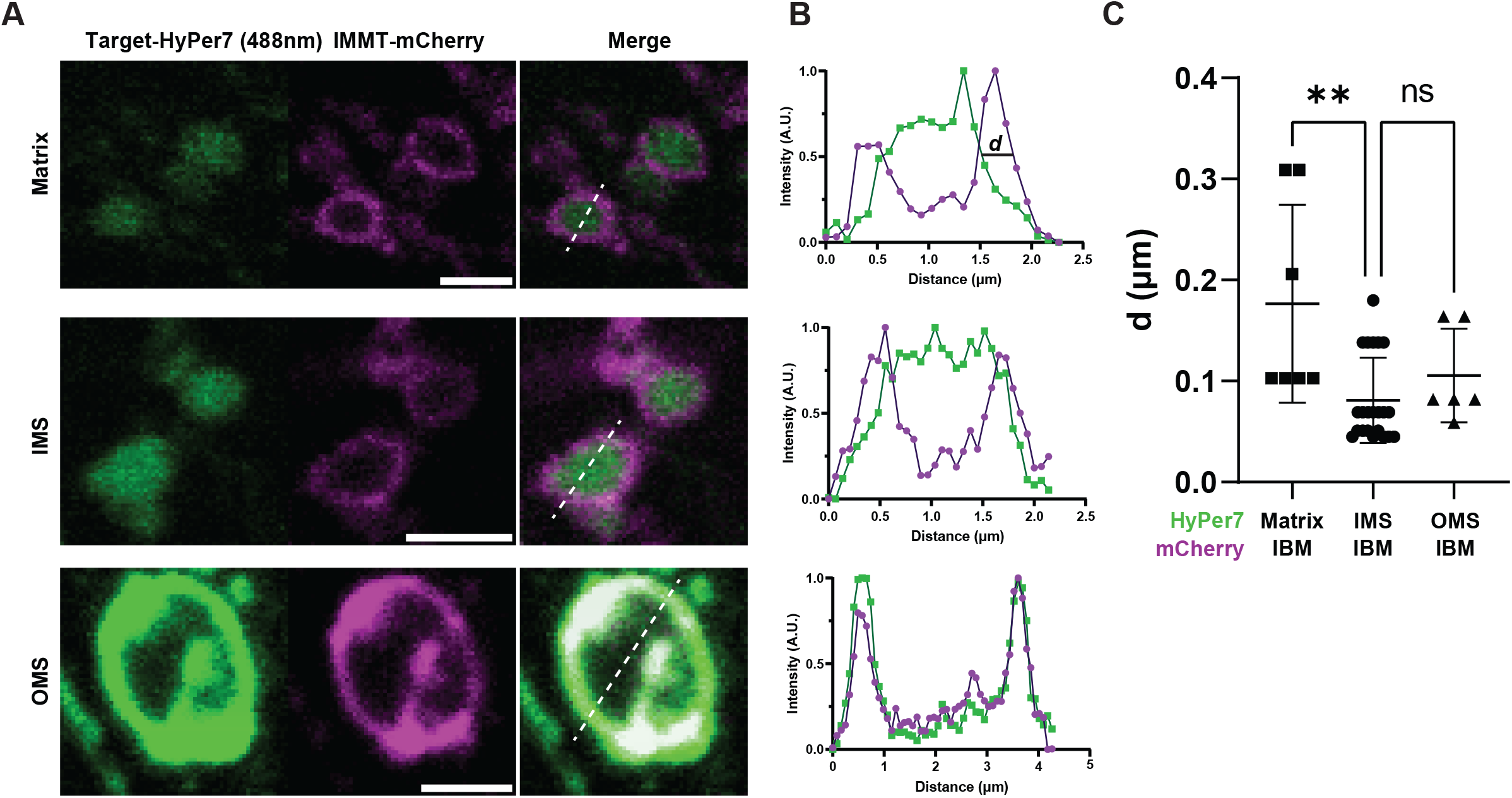
Single mitochondrial linescans of microdomain-targeted HyPer7. a) Representative image of mitochondrial linescans of matrix, IMS, and OMS-HyPer7 mitochondria in IMMT-mCherry expressing *C. elegans*. Scale bar represents 2 μm. b) Representative linescan data from A indicating distance metric between HyPer7 microdomain and the IBM (IMMT-mCherry). c) Comparison of distance metric between microdomain-specific HyPer7 and IBM-mCherry, mean ± SD, N = 6 – 22 mitochondria, one-way Kruskal-Wallis ANOVA with Dunn’s *post-hoc* correction. *p < 0.05, **p < 0.01.

**Extended Figure 3.**
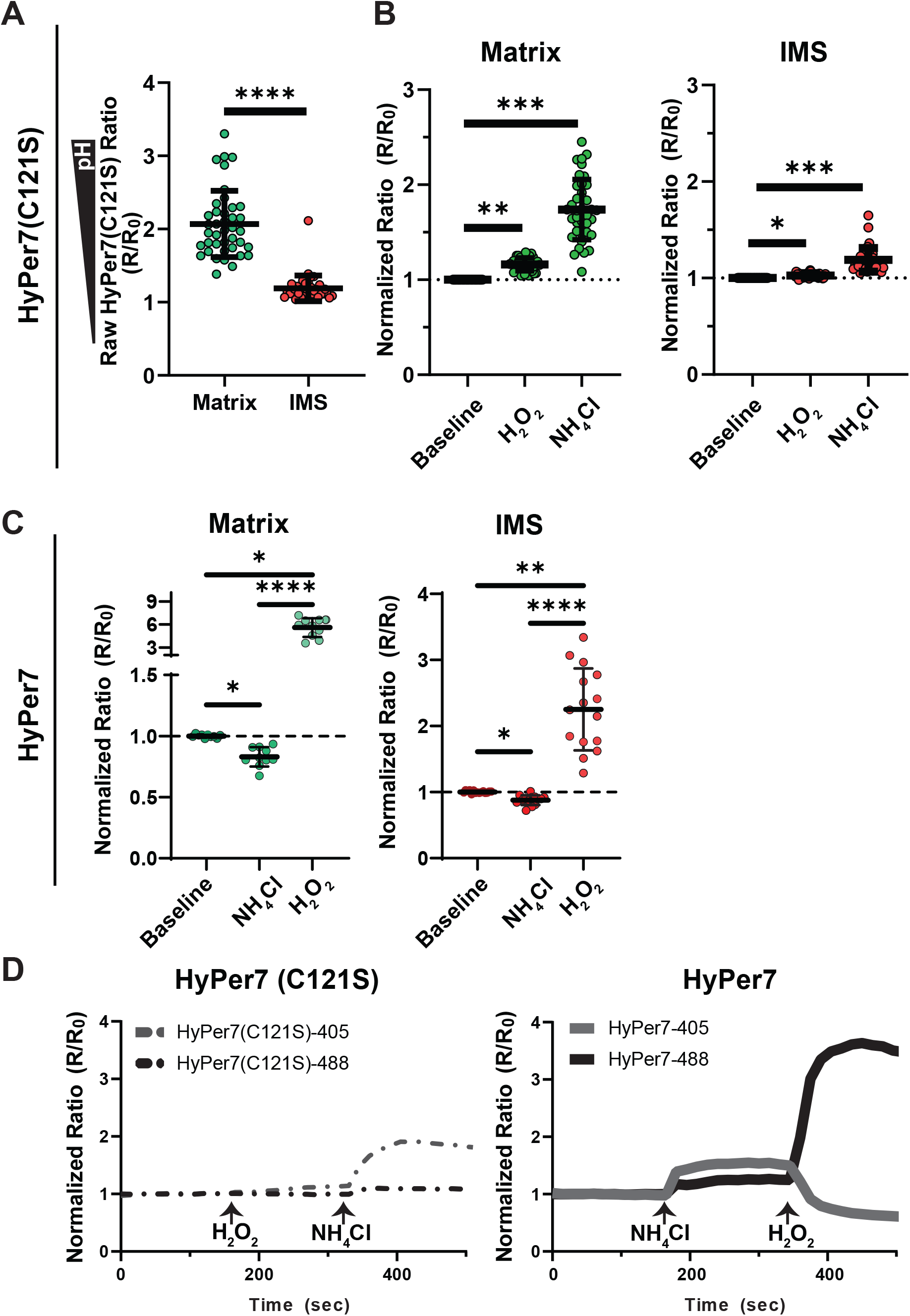
ROS and pH sensitivity of HyPer7 and ROS-insensitive HyPer7(C121S) variant. a) Raw HyPer7 ratio (488/405 nm excitation) of HyPer7(C121S) mutant targeted to mitochondrial matrix and IMS using two-tailed Mann-Whitney test, mean ± SD, N= 38 - 40 cells. b) Baseline-normalized HyPer7(C121S) mutant ratio (R/R_0_) of matrix- and IMS-targeted HyPer7 in response to saturating 100 μM H_2_O_2_ and 40 mM NH_4_Cl using Kruskal-Wallis one-way ANOVA with Dunn’s *post-hoc* correction, mean ± SD, N = 38 - 40 cells. c) As in b), but for HyPer7. N = 10 - 15 cells. d) Average traces of HyPer7(C121S) mutant and HyPer7 to H_2_O_2_ and NH_4_Cl treatment. *p < 0.05, **p < 0.01, ***p < 0.001.

**Extended Figure 4.**
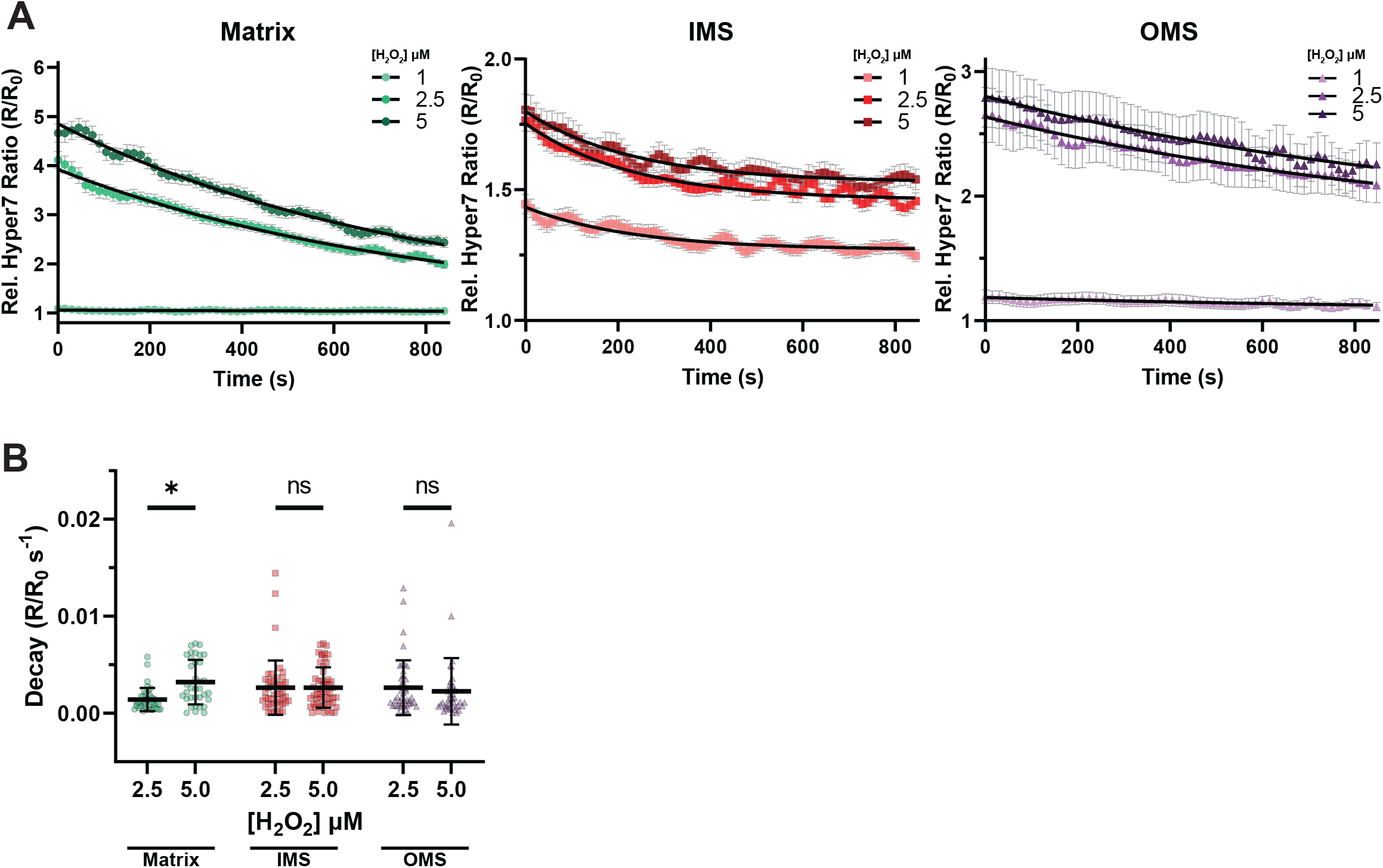
Microdomain-specific decay of HyPer7 following exogenous hydrogen peroxide. a) Relative HyPer7 ratio decay kinetics fit to nonlinear one-phase decay curves per microdomain after exposure to 1, 2.5, and 5 μM exogenous H_2_O_2_. N = 35 – 57 cells, mean ± SD. b) Statistical comparison of decay k constant of microdomain-specific HyPer7 ratio intensity over time after 1, 2.5, and 5 μM H_2_O_2_ treatment. Two-way ANOVA with Tukey post-hoc correction, N = 35 – 57 cells, mean ± SD. *p < 0.05

**Extended Figure 5.**
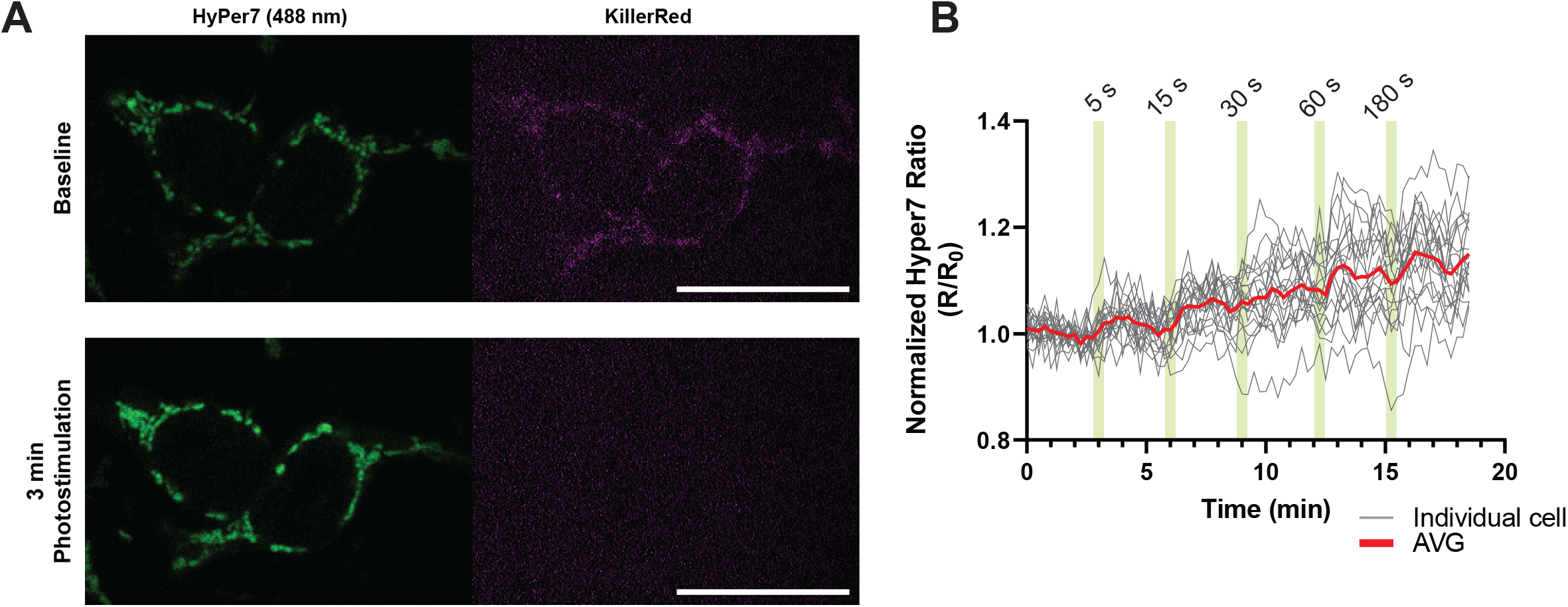
All-optical mitochondrial ROS generation and detection in live cells with kinetic parameters. a) Representative image of 488 nm excited matrix-HyPer7 and matrix-KR in cells before and after 180 sec of whole-cell photostimulation indicating complete photobleaching of KR. b) Individual cell (grey) and averaged (red) traces of dual excited ratiometric matrix-HyPer7 normalized to baseline (R/R_0_) of denoted durations of whole-cell KR photostimulation. Scalebar represents 25 μm.

**Extended Figure 6.**
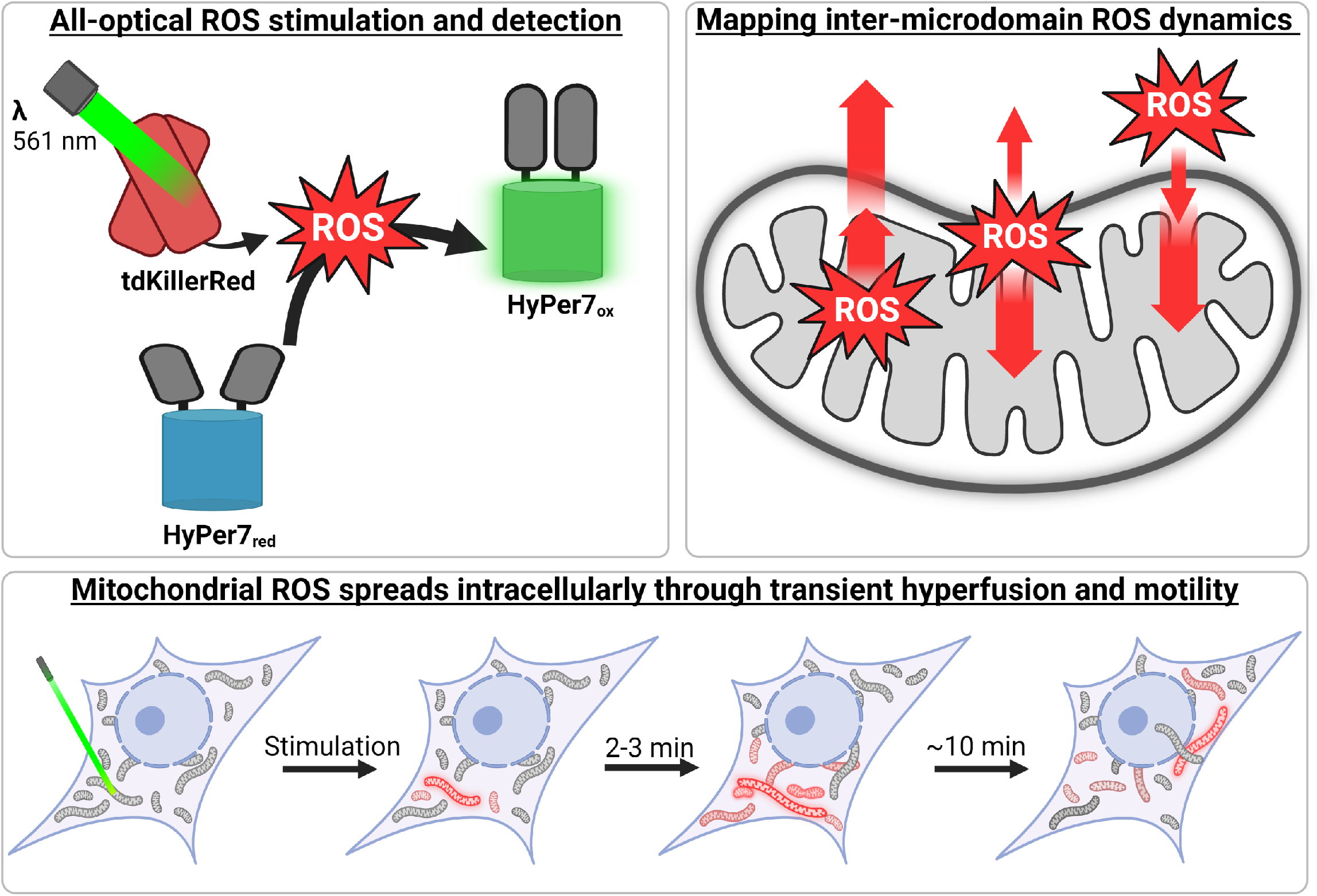
Graphical summary.

Supplemental Video 1: Matrix HyPer7 and KillerRed Photostimulation

Supplemental Video 2: No stimulation control Matrix HyPer7 and KillerRed

Supplemental Video 3: Matrix ROS induces motility of oxidized mitochondria

Supplemental Video 4: Matrix ROS induces transient hyperfusion of oxidized mitochondria

